# Insulin controls olfactory gain at the first central synapse by regulating periglomerular neuron excitability

**DOI:** 10.64898/2026.07.22.740090

**Authors:** Merve Oncul, Cédric Stefens, Emma Smith, Shreya Choudhuri, Beatrice Maria Filippi, Jamie Johnston

## Abstract

Sensory processing is dynamically tuned by internal state, yet how metabolic signals reshape the earliest stages of sensory circuits remains poorly understood. Here we identify a circuit mechanism by which satiety suppresses olfactory sensitivity at the first central synapse in the mouse olfactory bulb. Using a within-animal paradigm modelling fasted and glucose-induced sated states, we show that satiety impairs food-finding behaviour and reduces olfactory receptor neuron input to the olfactory bulb. Periglomerular (PG) cells, which co-express insulin receptors and the potassium channel Kv1.3, mediate this effect: insulin inhibits the low-voltage-activated Kv1.3 current in PG cells, increasing their spontaneous and odour-evoked activity. This heightened PG cell activity drives enhanced presynaptic inhibition of olfactory receptor neuron terminals, dampening sensory input before it reaches mitral cells. These findings establish insulin-dependent presynaptic inhibition of PG cells as a key locus of state-dependent sensory gain control.

## Introduction

The sensitivity of the olfactory system is not fixed; rather, it is continuously adjusted according to the animal’s internal metabolic state, linking the detection of food odours directly to the need for food. Olfaction plays a key role in stimulating food consumption and regulating energy metabolism across mammals (*1-4*), making the olfactory system a critical interface between metabolic state and feeding behaviour. Despite this, the neural circuit mechanisms by which metabolic signals modulate olfactory processing remain poorly understood.

The influence of metabolic state on olfactory sensitivity is well established at the behavioural level. Fasting enhances, and satiety reduces, olfactory detection in both rodents and humans (*5, 6*). Insulin is released postprandially in proportion to the glycaemic response and crosses the blood-brain barrier (*7*), making it a strong candidate signal for communicating satiety to the brain. These effects are not merely correlative: direct elevation of insulin within the olfactory bulb, either via intranasal delivery or local application, is sufficient to recapitulate the reduction in olfactory sensitivity associated with satiety (*8-12*), implicating insulin as a key mediator. Nevertheless, the specific circuit elements and cellular mechanisms through which insulin exerts these effects have not been identified.

The olfactory bulb expresses receptors for many signals involved in appetite regulation (*13, 14*) with the highest density of insulin receptors of any brain region (*15, 16*), making it a strong candidate site for metabolic modulation of olfactory processing. Within the glomerular layer, olfactory receptor neuron (ORN) terminals release glutamate onto mitral and tufted cells, while periglomerular (PG) cells provide both feedback and feedforward inhibition onto ORN terminals and mitral cells respectively (*17*). They can therefore act as a gate on the flow of sensory information (*18*). Insulin has previously been shown to act on mitral cells via inhibition of Kv1.3, increasing their intrinsic excitability (*19, 20*). This is paradoxical: if insulin increases mitral cell’s propensity to fire action potentials, olfactory sensitivity should increase with satiety, the opposite of what is observed behaviourally. This paradox has left the circuit basis of insulin’s effect unresolved. Periglomerular cells are well positioned to resolve this contradiction. Their dual role in feedback inhibition onto ORN terminals and feedforward inhibition onto mitral cells positions them as a gate on sensory input. Whether insulin acts on PG cells to suppress ORN input has not been tested.

Here we identify a circuit mechanism by which satiety suppresses olfactory sensitivity at its first central synapse. Using a within-animal paradigm combining behavioural, imaging, and electrophysiological approaches, we show that insulin acts on PG cells to gate olfactory receptor neuron input to the olfactory bulb, which can reduce olfactory function in the sated state.

## Results

### Glucose injection recapitulates satiety and reduces olfactory function

To examine how metabolic state modulates olfactory processing, we developed a within-animal paradigm in which fasted and sated conditions could be directly compared. Overnight food deprivation (16 h) produced a 10.6 ± 2.5% reduction in body weight (N = 45). A sated-like state was then induced by intraperitoneal glucose injection (2 g/kg), which reliably elevates both blood glucose and circulating insulin (*21*). Fasted mice injected with glucose consumed 52% less food during the first hour of re-feeding compared to saline-injected fasted controls (p = 0.00011, unpaired t-test; N = 6 and N = 5, respectively), consistent with previous work (*22*).

Satiety is associated with reduced olfactory sensitivity, an effect that can be recapitulated by insulin elevation within the olfactory bulb in both rodents and humans (*5, 9-1123*). To confirm whether our low- and high-energy paradigm altered olfactory ability, we performed a food finding test using mice that had been fed ethyl tiglate scented food over the preceding week. Following overnight fasting, and 15 mins after receiving an injection of saline or glucose, mice were placed in a cage with a scented food pellet buried beneath the bedding (Figure 1). Mice found the food in 66.6 ± 34.9 s after a saline injection, whereas the same animals took almost twice as long to locate the food following a glucose injection (Figure 1B, 127.6 ± 95.2 s; paired t-test, p = 0.029). The location of where mice dug during the search phase was markedly different between conditions (Figure 1A&C). For each mouse we measured the median distance from the buried food at which digging occurred. Saline-treated mice focused most of their digging almost directly over the food (1.8 ± 0.7 cm from the pellet), whereas after glucose the same mice dug substantially further away and far more variably (9.0 ± 7.1 cm; paired t-test, p = 0.004). These data indicate that our glucose-induced high-energy state likely impairs olfactory function, as it reduced the spatial precision with which mice directed their digging toward the buried food.

**Figure 1:**
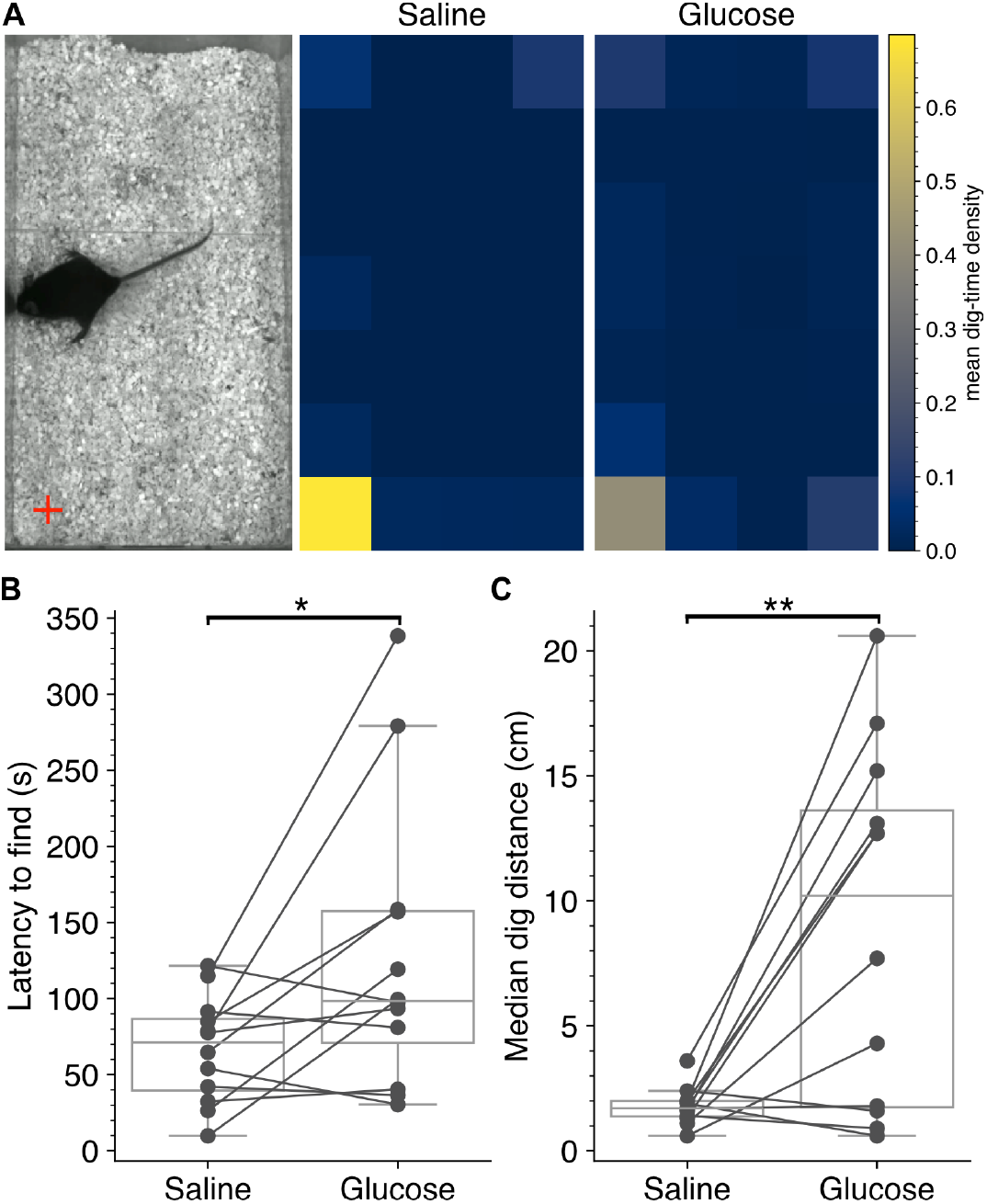
A glucose injection suppresses the performance of fasted mice in a food-finding test. **(A)** Left: a glucose-injected mouse digging during the food-finding test; location of food indicated by the red cross. Middle and right: heat maps showing the mean probability density of digging within the cage after injection of saline (middle) or glucose (right). Each square represents 17.7 cm^2^. **(B)** Latency to find the food was greater after a glucose injection (p = 0.029, paired t-test, N = 12). **(C)** Median dig distance was more variable and further from the food after glucose vs. saline (p = 0.004, paired t-test, N = 12).

### Satiety reduces the gain of olfactory input at the first central synapse

We have established that a glucose injection reduces olfactory ability (Figure 1). We next asked at what stage of olfactory processing this effect arises, beginning with the input to the olfactory bulb. PG cells are well positioned to regulate the ORN input to the olfactory bulb (Figure 2A), providing both tonic and activity-dependent feedback inhibition onto olfactory nerve terminals (*24*). To examine the functional coupling between ORN input and PG cell activity, we developed an approach that enables simultaneous measurement of both signals. We combined intrinsic optical imaging, which reports presynaptic ORN input to the olfactory bulb (*25, 26*), with fluorescence imaging of Ca^2+^ signals in the inhibitory PG neurons using anaesthetised VGAT-GCaMP6f mice (see Methods).

**Figure 2:**
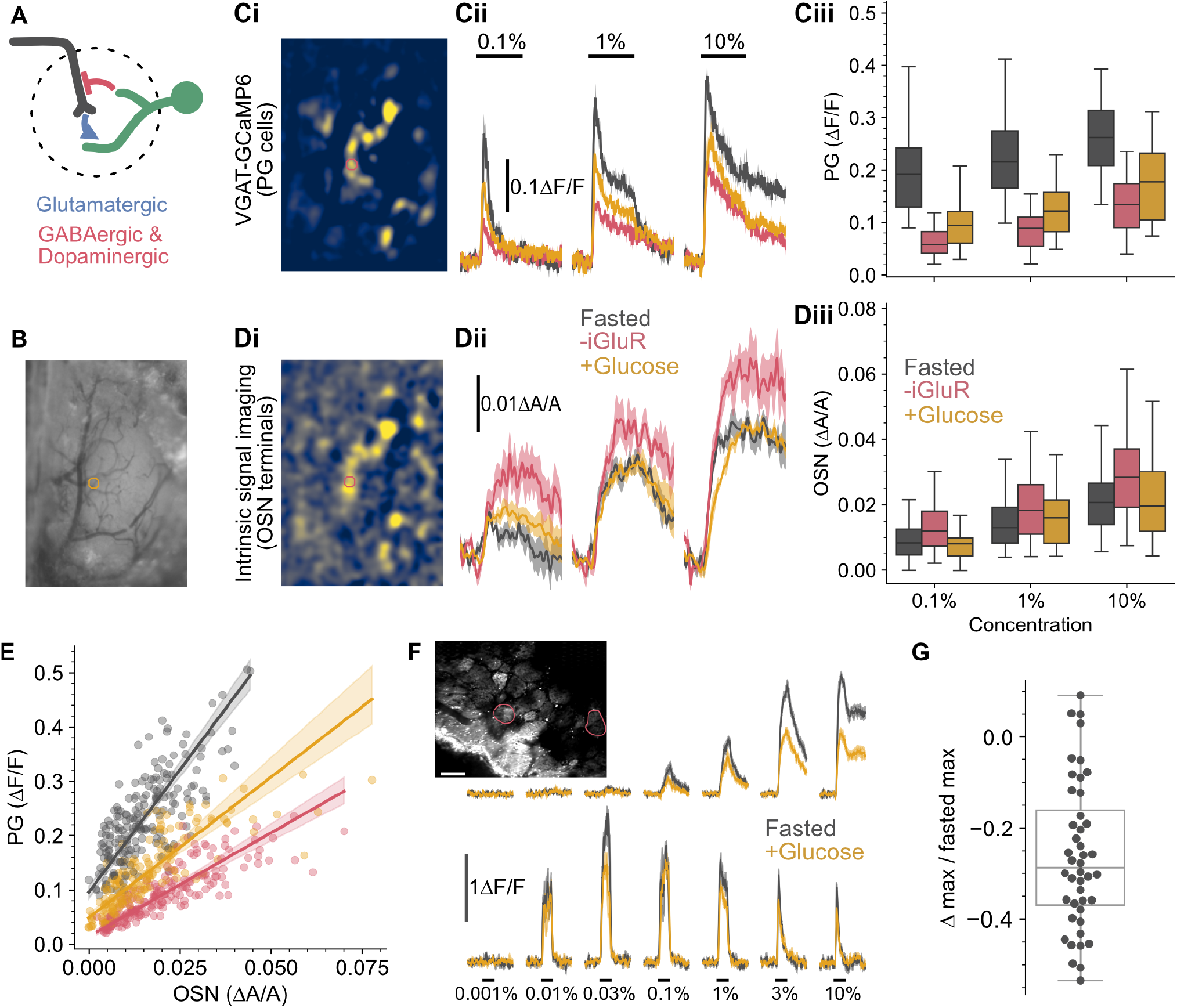
A glucose injection decreases olfactory nerve input yet increases glomerular inhibitory neuron activity. **(A)** The olfactory nerve input (black) forms a feedback loop with PG cells (green). **(B)** Wide-field imaging of the olfactory bulb, showing the field of view in C and D. **(Ci)** Band-pass filtered response map to 10% ethyl tiglate, showing VGAT-GCaMP6f glomerular activity (see Methods). **(Cii)** Example PG responses from the glomerulus indicated in Ci to ethyl tiglate, measured across 3 concentrations in the fasted state (black), in the presence of the ionotropic glutamate receptor antagonists NBQX and D-AP5 (red, “-iGluR”), and after a glucose injection (yellow). **(Ciii)** Summary data for all glomerular PG responses across concentration and state. Both concentration and state had a significant effect (p = 1.2 × 10^−29^ and p = 8.6 × 10^−83^, respectively; 2-way ANOVA), with no interaction. **(Di)** Band-pass filtered response map from intrinsic signal imaging, with the same glomeruli as in Ci indicated (see Methods). **(Dii)** Glomerular activity from ORN terminals for the glomerulus indicated in Ci and Di, using the same colour code as Cii. **(Diii)** Summary data for all glomerular ORN responses across concentration and state. Both concentration and state had a significant effect (p = 5.4 × 10^−40^ and p = 3.8 × 10^−8^, respectively; 2-way ANOVA), with no interaction. **(E)** Gain of PG cell responses to ORN input was reduced by ionotropic glutamate receptor antagonists (p = 4.0 × 10^−4^) and increased by glucose injection (p = 8.0 × 10^−4^; percentile bootstrap test, see Methods). **(F)** Two-photon imaging of ORN terminals in the olfactory bulb of an OMP-GCaMP6f mouse, showing the response of two glomeruli (indicated in inset) in the fasted and post-glucose state. **(G)** Fractional change in the maximal response of a glomerulus after a glucose injection. For C-E, data are from 62 glomeruli from 4 animals. Stimulus duration in Cii and Dii was 5 s, averaged across 5 trials. Stimulus duration in F was 3 s, averaged across 3 trials. Data shown as mean ± SEM. Scale bar in F = 100 µm.

In fasted mice, increasing odourant concentration produced a proportional increase in both ORN input and PG cell activity (Figure 2B–D, black traces). Consistent with the ORNs providing the primary excitatory drive to PG cells, ORN signal amplitude strongly predicted PG cell responses, accounting for 53% of the variance (Figure 2E, black; linear regression, R^2^ = 0.53, n = 186 glomeruli, N = 4 mice).

To directly perturb the excitatory drive onto PG cells and validate the sensitivity of our dual imaging approach, we applied the ionotropic glutamate receptor antagonists NBQX and D-AP5 to the surface of the olfactory bulb. This had 2 effects. As expected the reduced excitatory drive attenuated the amplitude of PG cell odour responses (Figure 2C black vs red, p = 2.7×10^-59^, Tukey post hoc); in contrast, the amplitude of the ORN responses was increased (Figure 2D black vs red, p = 3.8×10^-6^, Tukey post hoc), consistent with disinhibition due to reduced feedback inhibition from the smaller PG cell responses. These data confirmed that PG cells exist in a feedback loop with ORN input: blocking ionotropic glutamate receptors markedly attenuated PG cell odour responses even though the ORN terminal activity, which drives the PG cells was increased (Figure 2B–D, red traces). Accordingly, the relationship between ORN input and PG activity was attenuated (regression coefficient: 9.0, CI [8.2, 9.9] vs 3.8, CI [3.5, 4.1], p = 4.0 × 10^−4^), indicating reduced PG cell gain to ORN drive (Figure 2E, black and red). Because ORN input is the primary driver of PG activity, this slope provides a measure of PG cell sensitivity to ORN input, independent of ORN drive itself. This relationship therefore offers a way to isolate genuine changes in PG cell excitability from changes that simply reflect a shift in ORN input.

Having validated that our dual imaging approach can detect changes in PG cell gain to ORN input, we next asked whether our satiety paradigm alters this feedback loop. The same intraperitoneal glucose injection used in Figure 1 increased the amplitude of the PG cell response (Figure 2C, yellow, p = 3.5 × 10^−10^, Tukey post hoc), yet decreased the ORN input (Figure 2D, yellow, p = 2.7 × 10^−4^, Tukey post hoc). Critically, the gain of PG cell responses with respect to ORN input was also increased (3.8, CI [3.5, 4.1] vs 5.2, CI [4.7, 5.6], p = 8.0 × 10^−4^), confirming that this reflects a genuine increase in PG cell excitability rather than simply tracking the change in ORN drive. These data indicate that our satiety paradigm increases PG cell excitability and is associated with reduced olfactory nerve input to the olfactory bulb.

In a separate experiment, we imaged olfactory nerve terminals in OMP-GCaMP6f mice using two-photon microscopy. The same fasted/ glucose injection paradigm produced a 26.1 ± 16% reduction in the peak ORN response (p = 3.1 × 10^−4^, paired t-test, Figure 2F, G; n = 44, N = 4), consistent with the reduction in ORN input observed with intrinsic imaging above and corroborating that satiety reduces olfactory nerve input to the bulb.

### Insulin receptors and Kv1.3 are co-expressed in periglomerular cells

Elevated glucose results in increased circulating insulin (*21*), which we confirmed in our anaesthetised imaging paradigm: fasted insulin was 0.56 ± 0.013 ng/ml, rising to 0.83 ± 0.05 ng/ ml 40 minutes into the 55 minute post-glucose imaging protocol (N=6). Since insulin elevation within the olfactory bulb is sufficient to alter olfactory processing (*5, 9-1123*), we next asked whether the molecular targets of insulin signalling are expressed in PG cells. Insulin can modulate neural activity through two potassium channel targets: the voltage-gated channel Kv1.3 and the ATP-sensitive channel K_ATP_ (*19, 27, 28*). To determine whether both the insulin receptor and these downstream effectors are expressed in PG cells, we performed RNAscope multiplex fluorescence in situ hybridisation (Figure 3).

**Figure 3:**
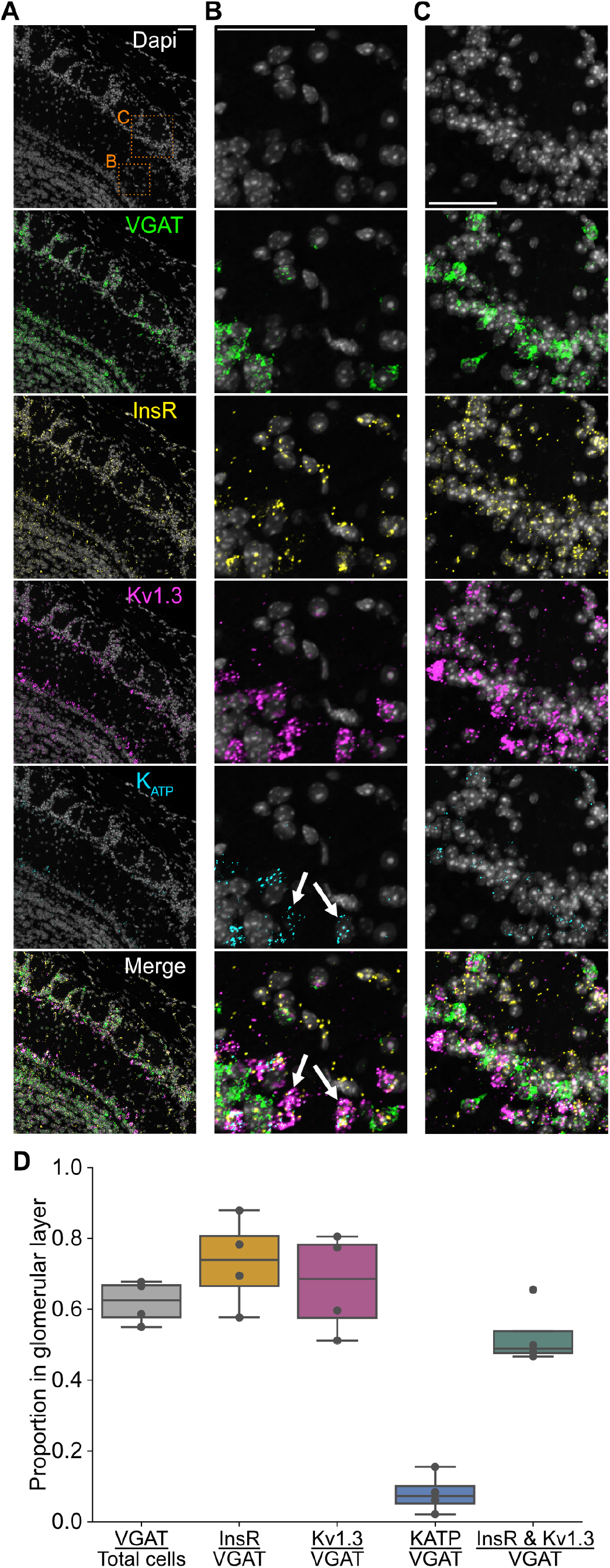
Insulin receptors and Kv1.3 channels are co-expressed by PG cells. **(A)** Overview image of an olfactory bulb section showing DAPI and probes for the insulin receptor (InsR), Kv1.3, and KATP, with a merged image; boxes indicate regions expanded in B and C. **(B)** Expanded view from A of the mitral cell layer; putative mitral cells indicated by white arrows. **(C)** Expanded view from A of the glomerular layer. **(D)** Proportions of probe-positive cells in the glomerular layer.

We restricted analysis to the glomerular layer, classifying cells as probe-positive if puncta were detected within the DAPI-defined nucleus (see Methods). Across four animals, we quantified 2,830 glomerular layer cells (708 ± 132 per mouse, N = 4). Of these, 62 ± 6% were VGAT positive (VGAT+), with the remainder likely comprising external tufted cells and glia. Among VGAT+ cells, Kv1.3 and the insulin receptor were co-expressed in 53 ± 9% (Figure 3C,D), indicating that the majority of PG cells express both targets. In contrast, K_ATP_ was expressed in only 8 ± 6% of VGAT+ cells, suggesting a more limited role for this channel in the PG cell population. Consistent with previous reports (*19*), putative mitral cells expressed both Kv1.3 and the insulin receptor, and also expressed K_ATP_ (^Figure^ 3B, white arrows).

### Insulin inhibits a low-voltage-activated potassium conductance in periglomerular cells

Kv1.3 is an insulin-sensitive low-voltage-activated K^+^ channel that opens in response to small depolarisations near the resting membrane potential, regulating action potential threshold and firing frequency (*29*). Given its expression in PG cells, we asked whether these neurons exhibit low-voltage-activated K^+^ currents that are sensitive to insulin.

To target PG and short-axon cells preferentially, we selected glomerular layer neurons with smaller somata for whole-cell voltage-clamp recordings. External tufted (ET) cells can be distinguished by their relatively low input resistances (∼200 MΩ; (*30, 31*), compared with short-axon and PG cells (>300 MΩ; (*30, 32-34*). Across 44 recorded glomerular layer neurons, input resistances ranged from 136 MΩ to 1.88 GΩ (Figure 4), with a strong bias toward higher values (median = 786 MΩ, IQR = 515-1,180 MΩ, n = 44, N = 24), consistent with successful targeting of the smaller-soma population.

**Figure 4:**
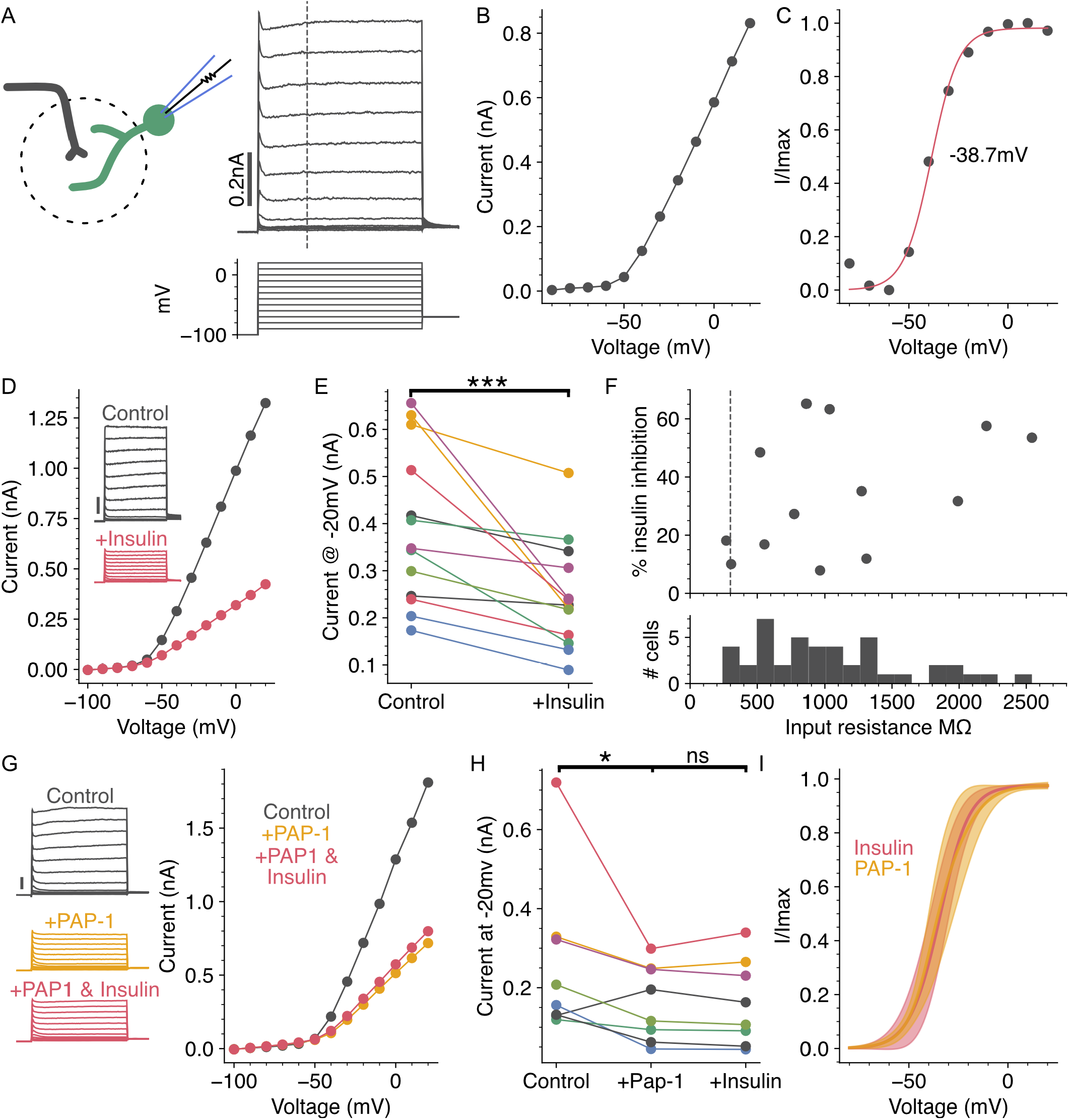
Insulin inhibits a voltage-gated potassium current active around resting membrane potential in PG cells. **(A)** Outward K+ currents were measured from glomerular neurons using 200 ms voltage steps between -90 and +20 mV, preceded by a -100 mV pre-pulse. **(B)** Current-voltage plot of the data in A, measured at the dashed line. **(C)** The data in B, corrected for the K+ driving force, normalised, and fit with a Boltzmann function (see Methods), giving a V0.5 of -38.7 mV. **(D)** A glomerular neuron before and after insulin application; inset scale bar, 0.2 nA. **(E)** Insulin caused a significant reduction in the K+ current measured at -20 mV (p = 0.002, paired t-test, n = 13, N = 11). **(F)** Percent inhibition of the current at -20 mV, plotted against input resistance for all 44 cells; dashed line at 300 MΩ. **(G)** A glomerular neuron before, after PAP-1, and after subsequent addition of insulin. **(H)** PAP-1 caused a significant reduction in the K+ current measured at -20 mV; subsequent application of insulin had no further effect (p = 0.047 and p = 0.64, respectively; corrected Wilcoxon signed-rank, n = 8, N = 7). **(I)** Mean activation curves of the insulin-sensitive and PAP-1-sensitive K+ currents. Only cells with >20% inhibition were used for the averages, shown ± 1 SD.

To characterise the voltage-gated K^+^ current profile, we evoked outward K^+^currents in whole-cell voltage-clamp in the presence of 1 µM TTX. Currents were elicited by a series of voltage steps from -90 to +20 mV preceded by a 750 ms pre-pulse to -100 mV to fully relieve channel inactivation (Figure 4A-B). The normalised conductance-voltage relationship, corrected for the K^+^driving force using the Goldman-Hodgkin-Katz current equation, was well fit by a Boltzmann function (Figure 4C). The median half-activation voltage was -31 mV (IQR -37 to -25 mV, n = 44, N = 24), indicating that the majority of recorded PG cells express low-voltage-activated K^+^ channels; only 9 of 44 cells had half-activation voltages more positive than -20 mV.

Insulin application caused a 34 ± 20% (SD) reduction in outward K^+^ current measured at -20 mV (p = 0.002, n = 13, N = 11; Figure 4D-E). All cells exhibiting greater than 20% insulin-induced current reduction had input resistances exceeding 500 MΩ (Figure 4F), consistent with a PG cell identity. Activation curves fitted to the insulin-sensitive current component yielded a half-activation voltage of -32 ± 2 mV (n = 13, N = 11), confirming that insulin preferentially inhibits a low-voltage-activated K^+^conductance.

To determine whether insulin’s effect is mediated by Kv1.3, we applied the Kv1.3 antagonist PAP-1, which reduced outward K^+^ current at -20 mV by 31 ± 35% in 8 cells (Figure 4G-H). Subsequent co-application of insulin in the continued presence of PAP-1 produced no further reduction in current (p = 0.64, Wilcoxon signed-rank), indicating that the insulin-sensitive current is fully occluded by Kv1.3 blockade (Figure 4G-H). The activation curve of the PAP-1-sensitive current closely matched that of the insulin-sensitive current recorded in the absence of PAP-1 (-34 ± 6 mV; Figure 4I). These data demonstrate that insulin inhibits a low-voltage-activated K^+^ conductance in PG cells through suppression of Kv1.3, consistent with the co-expression of the insulin receptor and Kv1.3 identified by RNAscope (Figure 3).

### Insulin increases spontaneous and evoked periglomerular cell activity

Inhibition of Kv1.3 by insulin would be expected to increase PG cell excitability (*29*), elevating both spontaneous activity and responses to olfactory nerve input. To test this prediction, we prepared acute slices from VGAT-GCaMP6f mice and used 2-photon calcium imaging to measure both olfactory nerve-evoked and spontaneous PG cell activity (Figure 5). To isolate postsynaptic changes in PG cell excitability from indirect effects of altered presynaptic release, experiments were conducted in the presence of the GABA_B_ receptor antagonist CGP 55845 (10 µM) and the D2 receptor antagonist sulpiride (100 µM). Under these conditions, insulin significantly increased olfactory nerve-evoked calcium responses (Figure 5B-C, Friedman test, p = 6.48 × 10^−22^), with significant increases at each individual stimulus intensity (Wilcoxon signed-rank test, corrected p = 1.66 × 10^−7^, 1.37 × 10^−12^, 1.28 × 10^−12^, 1.37 × 10^−12^, and 9.57 × 10^−9^, from weakest to strongest stimulus). Spontaneous activity, measured as the RMS of the ΔF/F during the inter-stimulus interval, was also significantly increased following insulin application (p = 3.48 × 10^−22^, Wilcoxon signed-rank test, n = 287 cells, N = 3 mice; Figure 5E).

**Figure 5:**
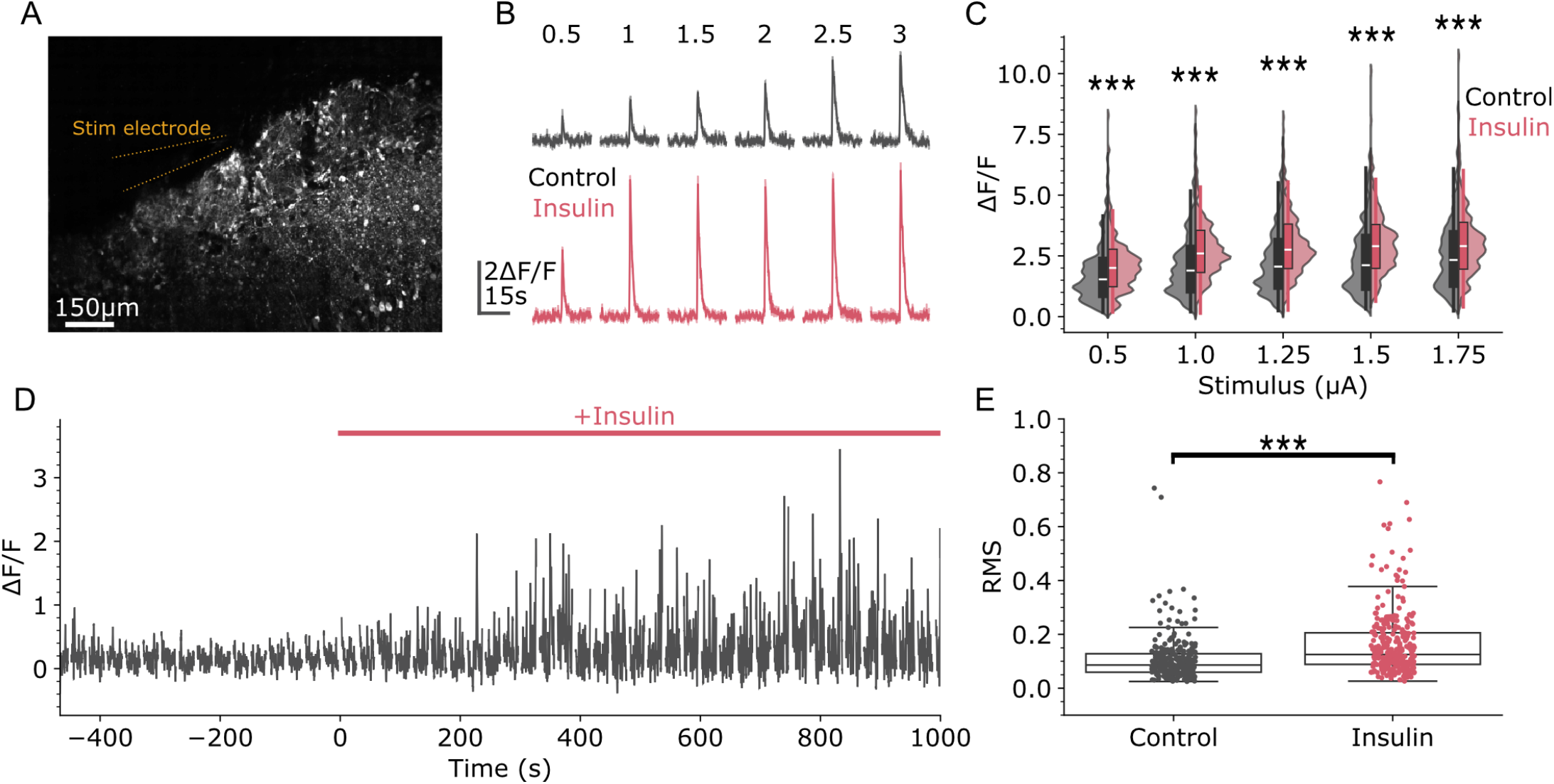
Insulin modulates PG cell activity. All recordings were conducted in the presence of 10 µM CGP 55845 and 100 µM sulpiride. **(A)** Field of view of PG cells in a slice from a VGAT-GCaMP6f mouse, with a stimulus electrode placed in the olfactory nerve layer. **(B)** Example olfactory nerve-evoked Ca^2+^ responses before (black) and after insulin (red); mean ± SEM of 3 repeats. **(C)** Insulin significantly increased evoked responses (p = 6.48 × 10^−22^, Friedman test), with significant increases at all stimulus intensities (p = 1.66 × 10^−7^, 1.37 × 10^−12^, 1.28 × 10^−12^, 1.37 × 10^−12^, and 9.57 × 10^−9^, weakest to strongest; corrected Wilcoxon signed-rank; n = 287, N = 3). **(D)** Example PG cell recording showing insulin’s effect on spontaneous activity; olfactory nerve stimuli have been blanked. **(E)** Insulin increased spontaneous PG cell activity, measured as RMS ΔF/F during inter-stimulus intervals (p = 3.48 × 10^−22^, Wilcoxon signed-rank; n = 287, N = 3).

### Insulin suppresses olfactory nerve terminal activity via presynaptic inhibition

Having established that insulin increases PG cell activity, we next asked whether this translates into enhanced feedback inhibition onto olfactory nerve terminals. To test this prediction, we repeated the imaging protocol in OMP-GCaMP6 mice, in which GCaMP6 is expressed selectively in olfactory sensory neurons, allowing direct measurement of presynaptic calcium transients as a proxy for olfactory nerve terminal activity (Figure 6).

**Figure 6:**
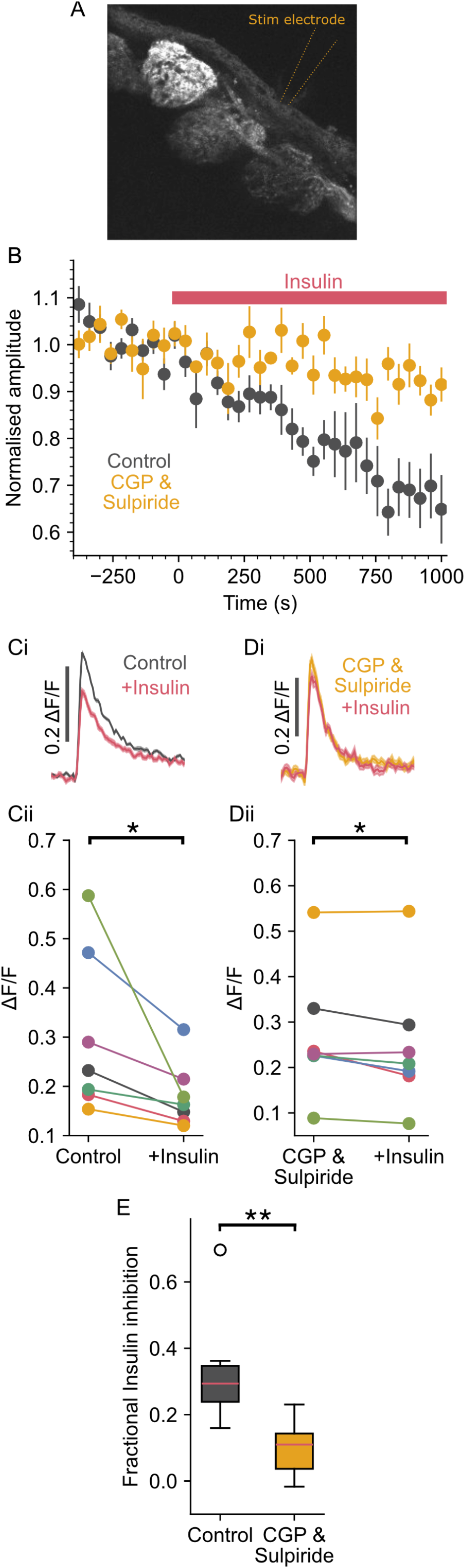
Insulin inhibits evoked calcium responses in ORN presynaptic terminals via feedback inhibition. **(A)** Field of view of glomeruli in a slice from an OMP-GCaMP6f mouse, with a stimulating electrode placed in the olfactory nerve layer. **(B)** Time course of insulin’s effect on olfactory nerve-evoked presynaptic Ca^2+^ transients, in control (black) and in the presence of 10 µM CGP 55845 & 100 µM sulpiride (yellow); responses normalised to the mean response prior to insulin application. **(Ci)** Example evoked glomerular ORN Ca^2+^ responses before (black) and after insulin (red); mean ± SEM of 20 stimuli. **(Cii)** Insulin reduced the maximum response from 0.30 ± 0.15 to 0.18 ± 0.06 ΔF/F (p = 0.016, N = 7). **(Di)** Example evoked glomerular ORN Ca^2+^ responses in the presence of CGP 55845 & sulpiride (yellow), before and after insulin (red); mean ± SEM of 20 stimuli. **(Dii)** In the presence of CGP 55845 & sulpiride, insulin reduced the maximum response from 0.27 ± 0.13 to 0.25 ± 0.14 ΔF/F (p = 0.041, N = 7). **(E)** Fractional inhibition by insulin was smaller in the presence of CGP 55845 and sulpiride (p = 0.008).

Insulin reduced olfactory nerve-evoked calcium transients in OMP-GCaMP6 mice by 33 ± 16% (Figure 6B, C & E, p = 0.016, Wilcoxon signed-rank, N = 7), consistent with increased presynaptic inhibition driven by the enhanced PG cell activity described in Figure 5. Critically, this effect was largely abolished when the same experiment was conducted in the presence of CGP 55845 (10 µM) and sulpiride (100 µM) (Figure 6B, D & E), confirming that the insulin-induced reduction in olfactory nerve input is mediated by presynaptic inhibition from PG cells. Even in the presence of these presynaptic blockers, insulin still produced a small reduction in the presynaptic response (9.7 ± 8.1%, p = 0.041, N = 7, Figure 6B, D), but this residual reduction was significantly smaller than the reduction seen without the blockers (33 ± 16%, p = 0.008, unpaired t-test, Figure 6E). Although a direct effect of insulin on ORN axons or presynaptic terminals cannot be excluded, the modest and gradual nature of this reduction, relative to the insulin-induced reduction seen without presynaptic blockers, suggests it more likely reflects gradual rundown of responses from the isolated terminals over the recording period. Together, these data support a model in which insulin acts postsynaptically on PG cells to suppress Kv1.3-mediated currents, increasing PG cell activity and thereby enhancing presynaptic inhibition of olfactory nerve terminals.

## Discussion

We identified a circuit mechanism by which satiety suppresses olfactory sensitivity at the first central synapse in the olfactory pathway. Using a within-animal paradigm combining behavioural, imaging, and electrophysiological approaches, we show that elevated insulin increases PG cell excitability by inhibiting Kv1.3-mediated potassium currents (Figures 3-5). This enhances presynaptic inhibition onto ORN terminals, reducing sensory drive to the olfactory bulb (Figures 2 & 6). This likely contributes to the reduction in olfactory sensitivity (*8-12*) and decreased ability to localise food (Figure 1). Critically, PG cells also provide feedforward inhibition onto mitral cells (*17, 18*), meaning that insulin-driven increases in PG cell activity will likely suppress mitral cell output through two convergent mechanisms: the reduced excitatory drive from ORN terminals, reported here, and direct postsynaptic inhibition of mitral cells.

Previous work established that insulin acts on mitral cells via Kv1.3 inhibition, increasing their intrinsic excitability (*19, 20, 35*). This is paradoxical given that, behaviourally, satiety reduces rather than enhances olfactory sensitivity and has left the circuit basis of insulin’s effect on olfactory processing unresolved. Our findings provide a resolution: although insulin increases the intrinsic excitability of both mitral and PG cells via Kv1.3 inhibition, the net effect on olfactory output is likely dominated by the PG cell arm of this signalling. Insulin-driven increases in PG cell activity reduce ORN glutamate release onto mitral cell dendrites through presynaptic inhibition, while simultaneously suppressing mitral cell activity through feedforward GABAergic inhibition. Indeed, spontaneous IPSCs recorded in mitral cells are increased by insulin in acute slices (*20*). These two convergent inhibitory drives are likely sufficient to override any increase in mitral cell intrinsic excitability, producing the net reduction in olfactory sensitivity observed behaviourally. This illustrates a broader principle: that the behavioural consequence of a neuromodulator cannot be predicted from its effect on a single cell type in isolation, but depends on how it acts across the circuit as a whole.

Our experimental paradigm uses an intraperitoneal glucose injection to model the sated state, which reliably elevates circulating insulin (*21*). We show that this decreases the ability to localise food (Figure 1) and decreases olfactory receptor neuron input to the olfactory bulb (Figure 2). To affect the olfactory circuitry any circulating factor must cross the blood-brain-barrier and enter the extracellular space. Glucose changes from ∼0.9 mM to ∼1.3 mM in the extracellular space of the olfactory bulb following a similar IP injection or between fasted and fed states (*36*). Although extracellular brain insulin is much harder to measure, early microdialysis studies found fasted to fed increase of insulin of over 200% (*37*), though the viability of such approaches has been questioned (*38*). Whole tissue extracts of olfactory bulb comparing insulin content in fasted vs fed states show over a 2-fold increase (*9*). Local elevation of insulin within the olfactory bulb also recapitulates satiety-induced decreases in olfactory sensitivity (*9, 23*). It is therefore likely that the glucose injection affects olfactory function and olfactory bulb input by acting through the parallel increase in insulin. Indeed our slice experiments, where glucose was held constant, show that insulin inhibits olfactory nerve input (Figure 6). This is due to insulin inhibiting Kv1.3 channels in periglomerular neurons (Figures 3&4) increasing their spontaneous and evoked responses (Figure 5). Nevertheless, we cannot rule out a potential contribution from glucose itself. Future experiments using selective insulin receptor blockade or local insulin dialysis will be needed to fully dissociate these contributions. It also remains an open question whether other metabolic signals that converge on the olfactory bulb, such as leptin and ghrelin, act through this same presynaptic gate or engage distinct circuit elements.

Our findings may have relevance to conditions in which insulin signalling is chronically disrupted. Olfactory deficits are well documented in type 1 and type 2 diabetes, as well as in obesity and insulin resistance (*39-42*), yet their mechanistic basis is poorly understood. The circuit mechanism we identify, in which insulin acts on PG cells to gate ORN input via Kv1.3, provides a candidate cellular pathway through which disrupted insulin signalling could impair olfactory function in metabolic disease. Intranasal insulin delivery, which targets the olfactory bulb directly, has been shown to reduce olfactory sensitivity in healthy humans (*10,11*), consistent with the circuit mechanism we describe. These observations raise the possibility that the olfactory deficits associated with metabolic disease reflect, at least in part, dysregulation of insulin-dependent gain control at the first central olfactory synapse.

Together, our findings establish PG cell-mediated presynaptic inhibition as a key locus of metabolic gain control in the olfactory system, and provide a circuit-level framework for understanding how internal state shapes sensory processing at its earliest central stage.

## Methods

### Animals

Animal handling and experimentation was carried out according to UK Home Office guidelines and the requirements of the United Kingdom Animals (Scientific Procedures) Act 1986 and the University of Leeds animal welfare ethical review board. Mice were housed under a 12:12 h light/dark cycle with free access to food and water. All efforts were made to minimise animal suffering and the number of animals used. VGAT-Cre (stock 028,862, B6J.129S6(FVB) Slc32a1<tm2(cre)), and OMP-Cre mice (B6;129P2(Cg)-Omp<tm4(cre)Mom>/ MomTyagRbrc (RBRC02138)) were crossed with floxed-GCaMP6f mice (GCaMP6f.flox, stock 028,865, B6J.CgGt(ROSA)26Sor < tm95.1 (CAGGaMP6f)), to generate VGATxGCaMP6f mice, and OMPxGCaMP6f mice, respectively. The OMP-Cre mouse line was originally obtained from RIKEN BioResource Research Center (Ibaraki, Japan), with permission from P. Mombaerts, the original developer of the OMP-cre line (*43*). VGAT-cre and floxed-GCaMP6f were originally from Jackson Laboratory (Maine, USA). Wild-type mice (C57Bl6J) were obtained from Charles River Laboratories. All mouse lines were maintained in house. Consistent with the NC3Rs guidelines (https://www.nc3rs.org.uk/who-we-are/3rs), both males and females were used in this study.

#### Food consumption

After overnight fast, mice were given either a saline or glucose (2g/kg) intraperitoneal injection. 10 mins later they were given food and consumption was measured after 1 hr.

### Food Finding Test

The diets of 12 mice (6 of each sex) were supplemented with ethyl tiglate (0.01% w/v) for 7 days as in (*44*). Mice were then fasted overnight and received an injection of either saline of 2g/kg glucose (50/50 split). 15 mins after the injection they took part in a food finding test: a single odourised food pellet was buried in one corner of a cage filled with ∼4 cm of bedding, and the mouse was introduced and allowed to search for 10 mins or until it uncovered the pellet. 4 days later the food finding test was repeated with each mouse receiving the opposite injection. The pellet location was varied across mice and trials. Behaviour was filmed from above with at Basler ace (acA1300-200um) monochrome camera at 33 frames/s. Analysis was restricted to the search phase up until the food pellet became visible. Frames when the mouse exhibited digging were manually labelled and the location of the head was tracked using DeepLabCut (*45*) using the SuperAnimal-TopViewMouse model (*46*). For each animal, the head point at every detected digging frame within the search window was normalised to the cage rectangle, and the coordinate frame of each animal was reflected horizontally and/or vertically so that the food corner was placed in the bottom left corner of all maps. Normalised positions were binned into a grid whose aspect ratio matched the cage, the per-animal map was normalised to unit sum (dig-time density), and maps were averaged across animals within each condition; bin values therefore represent the mean fraction of digging time at each cage location, with square cells in real space. Distance to food was computed per detected dig as the median Euclidean distance between the animal’s body point and the manually marked food location.

### Odour stimulation

Ethyl tiglate was obtained from Sigma-Aldrich Liquid dilutions of Ethyl tiglate (Sigma-Aldrich) were prepared by serial dilutions in oil either (Sigma-Aldrich, 69794) or (Spectrum Chemical, C3465) within ∼1 week of experiments. Diluted odourants were delivered in vapour phase in synthetic medical air using either an 8- or 16-channel olfactometer (Aurora Scientific, 206A or 220A, respectively). Total flow rates from the olfactometers were kept constant at 1000 sccm (standard cubic centimetre per minute). The output tubing of the olfactometer was positioned 1 to 2 cm in front of the mouse’s nose. Inter-stimulus intervals were between 20 and 60 s for higher concentrations to minimise any adaptation. All odour concentrations are reported as % saturated vapour (sv).

### Optical multimodal imaging

VGAT-GCaMP6 mice were anaesthetised with urethane (1.5 g/kg) and body temperature was maintained at 37°C throughout. Animals were secured with a custom-made head bar and a craniotomy covering the olfactory bulb was performed. The exposed bulb was covered with 2% low-melting-point agarose in artificial cerebrospinal fluid and sealed with a glass coverslip, affixed with silicone rubber.

The olfactory bulb was imaged using a custom-built wide-field microscope comprising an Andor Zyla 5.5 camera (Oxford Instruments) mounted on an Olympus trinocular, with a long-pass filter (FELH0500, Thorlabs) and an inverted Canon EF 50 mm f/1.8 lens serving as the objective. Illumination was provided by collimated LEDs (Thorlabs) positioned directly over the cranial window: a green LED (M530L3) for imaging blood vessels and surface structure, a blue LED (M470L3) with a 457.9/10 nm clean-up filter for GCaMP6 excitation, and a red LED (M810L3) for intrinsic signal imaging. LEDs were driven by Cyclops LED drivers (Open Ephys), with the 810 nm LED under optical feedback control. Experimental control was via custom-written software (*47*). The three imaging modalities were acquired on successive frames, with each channel sampled at ∼18 Hz; the structural channel was used to register the other two channels to the first frame.

To identify glomeruli, response images were generated by averaging frames across the stimulus period and subtracting the pre-stimulus baseline. These response maps were spatially band-pass filtered (14-110 µm) to isolate glomerular signals from diffuse background activity, as described previously(*25*), a step that has been shown to substantially increase the spatial correlation between optically identified regions and histologically defined glomeruli (*25, 26*)(Meister & Bonhoeffer, 2001; Vincis et al., 2012). Responding glomeruli were then identified automatically in the band-pass-filtered maps using skimage’s Laplacian-of-Gaussian blob detector. This was performed independently on the intrinsic signal and GCaMP6 channels to generate a composite glomerular mask.

For each identified glomerulus, the time course was extracted from the raw image data together with that of a surrounding annulus (147 µm diameter), which was blanked wherever it overlapped a neighbouring glomerulus. The annular time course was subtracted from the glomerular signal (weighting of 0.7) to remove diffuse background contamination. Signals were then expressed as ΔF/F for GCaMP6 and as fractional change in absorbance relative to baseline (ΔA/A) for intrinsic signal imaging.

Glutamate receptor blockers were applied to the olfactory bulb by briefly removing the coverslip and replacing the agarose with agarose containing 1 mM D-AP5 and 0.1 mM NBQX, before resealing with a fresh coverslip.

### In vivo 2-photon calcium imaging

Male and female mice, ages between 60-90 days were anaesthetised with urethane (1.5 g/kg), and body temperature was maintained at 37°C. Animals were secured with a custom-made head bar, and a craniotomy covering the right hemisphere of the olfactory bulb was performed. The exposed bulb was covered with 2% low–melting point agarose in artificial cerebrospinal fluid, and a 3-mm glass coverslip (Biochrom) was affixed with dental cement. Silicone rubber (Body Double Fast Set) was applied to the skull surrounding the cranial window to create a well for the water dipping objective of the microscope. GCaMP6f fluorescence was imaged with a custom-built microscope, excited at 940 nm using a pulsed Mai Tai eHP DeepSee TI:Sapphire laser system (SpectraPhysics). A resonant-galvo mirror assembly (Sutter Instruments) scanned the beam through a 16× water-dipping objective (N16XLWD-PF, numerical aperture: 0.8, Nikon). Fluorescence was detected using GaAsP photomultiplier tubes and appropriate filters and dichroic mirrors. Images were acquired at 30 to 120 Hz, using ScanImage software (*48*).

### RNAscope Multiplex Assay

2 male and 2 female C57BL/6J mice were anaesthetised with pentobarbitone and fixed by transcardial perfusion of PBS followed by 4% PFA, the brains removed and fixed in PFA for 24 hrs. The fixed brains were then cryoprotected in 30% sucrose and embedded and flash frozen in OCT media. The olfactory bulbs were cryosectioned coronally at 12 µm. RNAscope was performed as per ACDbio manufacturers protocol. Probes were purchased (Bio-techne) for *Insr, KATP, Kv1*.*3*, and Vgat, and fluorophores Opal 480, Opal 540 (Akoya bioscience), and TSA Vivid 650, TSA 570 Vivid (Bio-techne) were assigned to each channel, respectively.

Images were aquired using Zeiss LSM880 + Airyscan Upright Confocal Microscope, (Axio Imager.Z2 microscope), 20X magnification (Plan-Apochromat 20x/0.8 objective), and Zeiss ZEN (black edition) software. For each image, a lambda scan was collected using lasers 458, 514, 561,633 along with a separate DAPI image. For analysis, normalised actual spectral data was obtained for each fluorophore. Each image was thresholded then probes were demixed by fitting a model incorporating the spectra for each pixel.

### Acute slice preparation

For electrophysiological recordings, P21-P60 female and male mice were used. Mice were deeply anaesthetized with isoflurane and decapitated. Coronal brain slices (300 µm thickness) containing the olfactory bulb were sectioned using a compresstome (Precissionary Instruments). Slices were prepared in chilled NMDG solution containing (in mM): 93 NMDG, 2.5 KCl, 1.2 NaH2PO4, 30 NaHCO3, 20 HEPES, 25 Glucose, 5 sodium ascorbate, 2 Thiourea, 3 Sodium pyruvate, 10 MgSO4.7H2O, 0.5 CaCl2.2H2O aerated with 95% O_2_/5% CO_2_ (pH is adjusted to 7.4 with HCl). Slices were transferred to an incubation chamber containing artificial cerebrospinal fluid (aCSF) containing (in mM): 125 NaCl, 4 KCl, 25 NaHCO3, 2 CaCl2, 1.25 NaH2PO4, 1 MgCl2, 5.5 Glucose, aerated with 95% O_2_/5% CO_2_. Slices were incubated at 32-35 °C for 10 min and then maintained at room temperature.

### Electrophysiology

Acute olfactory bulb slices were transferred to the recording chamber of a patch-clamp rig and perfused with aCSF at ∼35°C using a peristaltic pump and inline heater. Neurons in the glomerular layer were patched using pipettes with 4-7 MΩ tip resistances in the bath. The recordings were obtained using pipettes (Harvard Apparatus) made from borosilicate glass capillaries. Whole cell voltage-clamp recordings were performed using an Multiclamp 700B (Molecular Devices), signals where digitised using an NI-6356 A-D converter (National Instruments) controlled by Neuromatic software, which runs in Igor Pro (Wavemetrics). For voltage clamp recordings, a potassium gluconate based internal solution was used. Pipette solution contained (in mM): 135 K-gluconate, 10 KCl, 10 HEPES, 10 MgCl2, 2 Na2-ATP, 2 Na2-GTP (pH 7.3 with KOH; 280-285 mOsm). Neurons first received a 750 ms pre-pulse to -100 mV, to remove any inactivation and were then depolarised from -90 mV to a final potential of +20 mV, in 10 mV steps. Input resistance was measured using the 10 mV step between -80 and -70 mV. Activation parameters were determined by a Boltzmann function of the form I = I max/(1 + exp(V− V 1 /2 /k)) with variables I max, V 1/2 and k (the slope factor), with the non-linearity of the single channel current corrected using a modified GHK equation of the form i = α(F^2^V/RT) × {[K^+^]in − [K^+^]out·exp(−FV/RT)} / {1 − exp(−FV/RT)}, where the terms have their usual meaning and α is a normalisation factor (*49*). All voltages were corrected for a 13mV liquid junction potential.

### In vitro 2-photon calcium imaging

Acute olfactory bulb slices were transferred to the recording chamber, of a custom built 2-photon laser scanning microscope and perfused with aCSF at ∼35°C using a peristaltic pump and inline heater. GCaMP6f fluorescence was excited as described above. To electrically stimulate the olfactory nerve layer, we used a broken-tip patch electrode (5-10 µm diameter) filled with 0.9% and a constant current stimulator. Inter stimulus intervals were 30s.

### Pharmacological agents

Pharmacological tests were conducted by bath application of the following drugs with the indicated final concentrations in aCSF solution: TTX (1 µM), insulin (172 nM), PAP-1 (100 nM), CGP 55845 (10 µM) and sulpiride (100 µM).

### Quantification and statistical analysis

All data were tested for normality using the Shapiro-Wilk test; mean ± standard deviation and parametric tests are reported for normally distributed data, and median and interquartile range (IQR) and non-parametric tests are reported otherwise. Where appropriate, p-values were corrected for multiple comparisons using the Holm-Bonferroni method. Regression coefficients are reported with their confidence intervals.

## Acknowledgements

We would like to thank the University of Leeds CBS staff. **Funding**: This work was funded by grants from the MRC (MR/V003747/1) and BBSRC (BB/ Z51679X/1) and by a studentship funded by the University of Leeds. **Author contributions**: Writing - original draft: M.O., and J.J. Conceptualisation: C. S. and J.J. Investigation: M.O., C.S., E.S, S.C., Writing - review and editing: M.O., C.S., E.S, S.C., B.M.F., and J.J. Methodology: C.S., and J.J. Resources: J.J. Funding acquisition: J.J. & B.M.F. Data curation: C.S., M.O. and J.J. Validation: M.C., Z.Z., and J.J. Supervision: B.M.F., and J.J., Formal analysis: M.O., C.S. and J.J. Software: C.S., and J.J. Project administration: J.J. Competing interests: The authors declare that they have no competing interests.

